# Infanticide in a colonial cooperative bird is not associated with direct reproductive benefits

**DOI:** 10.1101/2025.05.27.656097

**Authors:** Rita Covas, Liliana R. Silva, André C. Ferreira, Pietro B. D’Amelio, Rita Fortuna, Delphine Duval, Matthieu Paquet, Claire Doutrelant

## Abstract

Infanticide, the killing of dependent young by adult conspecifics, is an extreme form of conflict that remains poorly understood. Previous studies suggest that it is associated with reproductive benefits, either by making breeders newly available for reproduction or by increasing offspring care and reducing competition for resources. In addition, infanticide may result from a reproductive trade-off, where parents kill their offspring if environmental conditions compromise their reproductive value. Here, we combine video, genetic and individual-level reproductive data collected over 12 years to describe infanticide in a highly social, cooperative bird, the sociable weaver *Philetairus socius* and to establish whether direct reproductive benefits or life-history trade-offs are possible motivators for this behaviour. We documented 50 cases of infanticide and identified 27 aggressors with known histories. These were mostly born outside the colonies where they committed infanticide (63%) and there was a bias towards younger individuals. Aggressors were usually not part of the victim’s breeding group (74%), but in two cases infanticide was committed by the breeding female, and in five by helpers. In most cases, aggressors were not reproducing at the time of the infanticide, were not recorded mating with any of the breeders or helpers from the attacked nest, nor occupied the nest from where the chicks were evicted. Only 13 of the known aggressors were confirmed as breeders after the attacks, and these mostly reproduced at the colony where the attacks were committed. Overall, we found no evidence for direct reproductive benefits of infanticide in sociable weavers. However, the younger age of aggressors, lack of genetic relatedness to the victims and future breeding at the colony suggest that competition between breeding groups, and hence indirect reproductive benefits, could provide motivation for infanticide in this population. Additional work over longer periods is needed to identify these possible indirect fitness benefits of infanticide.

## INTRODUCTION

Infanticide is an extreme form of conflict where dependent young are killed by adult conspecifics. It has been described in a wide range of taxa such as insects (Ratnieks, Foster and Wenseleers, 2006), fish (Jindal *et al*., 2017), mammals (Lukas and Huchard, 2014, 2019), reptiles (O’Connor and Shine, 2004) and birds (Heinsohn *et al*., 2011; Riehl, 2016; Cheng *et al*., 2019). From an evolutionary perspective, infanticide is intriguing because, in addition to the extreme cost for the victims, the costs for the aggressors can also be important. For example, studies in mammals found that infanticide was associated with an increase in aggressive behaviours between adults (both as a way of preventing infanticide and retaliation) and social exclusion of the individuals that could potentially commit infanticide in the future (see reviews in Agrell et al., 1998; Ebensperger, 1998). Hence, for infanticide to evolve and be maintained in any population, the fitness benefits to the actors should be more important than the costs incurred.

The benefits of infanticide appear to be most often associated with direct reproductive benefits, either through access to a mate or competition for resources for developing offspring. In mammals, comparative work has shown a link between infanticide and reproductive benefits that differ among the sexes (Lukas and Huchard, 2014, 2019). For males, killing young brings their mothers into earlier oestrous, which can make them available to reproduce with that male (the sexual selection hypothesis; (Hrdy, 1979) see also (Ebensperger, 1998; Lukas and Huchard, 2014). For females, infanticide appears to be associated with competition for the resources available for reproduction and the developing offspring (resource competition hypothesis; Hrdy, 1979; Lukas and Huchard, 2019), but the evolutionary determinants seem to be more complex than for males (Lukas and Huchard, 2019). The spatial proximity of other females can increase competition for food or the parental care provided to the offspring by other group members, but also for space (nest or dens; (Lukas and Huchard, 2019; Brown *et al*., 2021). Evidence for these different forms of resource competition has also been found in other taxonomic groups. For example, in the greater ani *Crotophaga major*, more than one female lay eggs in one nest and females can toss eggs or kill hatchlings of other females, leading to reduced starvation of their own chicks (Riehl, 2016). In green-rumped parrotlet *Forpus passerines*, where cavities to breed are in short supply, non-breeding pairs (or individuals) can commit infanticide in an attempt to gain access to the cavity where the young were developing (or to mate with one of the parents; Beissinger and Berg, 2024). Reproductive benefits can also arise indirectly, through kin selection or through competition among matrilines or clans for access to future reproduction, to higher social status (which is necessary for achieving reproductive status) or reduced competition for future offspring (Rubenstein, 2012; Lukas and Huchard, 2019; Brown *et al*., 2021). Finally, in social insects, workers caring for the developing larvae can identify those that are not their sisters and selectively kill them (Ratnieks, Foster and Wenseleers, 2006), providing an example where infanticide is favoured by indirect fitness benefits through spiteful behaviour (see also West and Gardner, 2010).

Life history trade-offs might also explain cases where infanticide is committed by parents. For example, in grey-capped social weavers *Pseudonigritta arnaudi,* there is evidence of infanticide when parents suffered partial predation in the nest (Cheng *et al*., 2019). In these cases, the parents ejected the whole clutch and started a new clutch instead of raising the remaining nestlings, as the cost of raising a partial clutch was unlikely to overcome the benefit (Cheng *et al*., 2019). A similar life-history trade-off was suggested in eclectus parrots *Eclectus roratus,* where killing male offspring appeared advantageous for parents in poorer-quality nest hollows, as broods with a single female offspring were found to have higher reproductive value than broods in which the female had a younger brother (Heinsohn *et al*., 2011). Further (albeit anecdotal) support for this hypothesis comes from a mammal, the golden takin *Budorcas taxicolor bedfordi*, where mothers were recorded to kill their newborns when they failed to stand up within the usual time (Zhao *et al*., 2020).

Hence, the benefits of infanticide appear to vary according to sex, reproductive status of the aggressors and reproductive costs and benefits. However, infanticide events are infrequent and can take place rapidly, which means that they are difficult to document. Therefore, infanticide and its evolutionary drivers remain poorly understood, in particular in non-mammalian animals.

Here, we use video recordings collected over 12 years of breeding monitoring to identify infanticide (Fig.1) in a highly social, colonial cooperative bird, the sociable weaver *Philetairus socius*. Using our individual-based long-term data, we describe the identity, individual and social attributes of the aggressors: sex, age, colony of origin and resident/immigrant status, and breeding and helping activity. Based on these data, we examine two types of hypotheses that could explain the adaptive significance of infanticide in this species, i) reproductive benefits arising either from sexual selection (access to a mate) or competition for resources (breeding chamber or food for developing offspring) and ii) life-history trade-offs.

**Figure 1:**
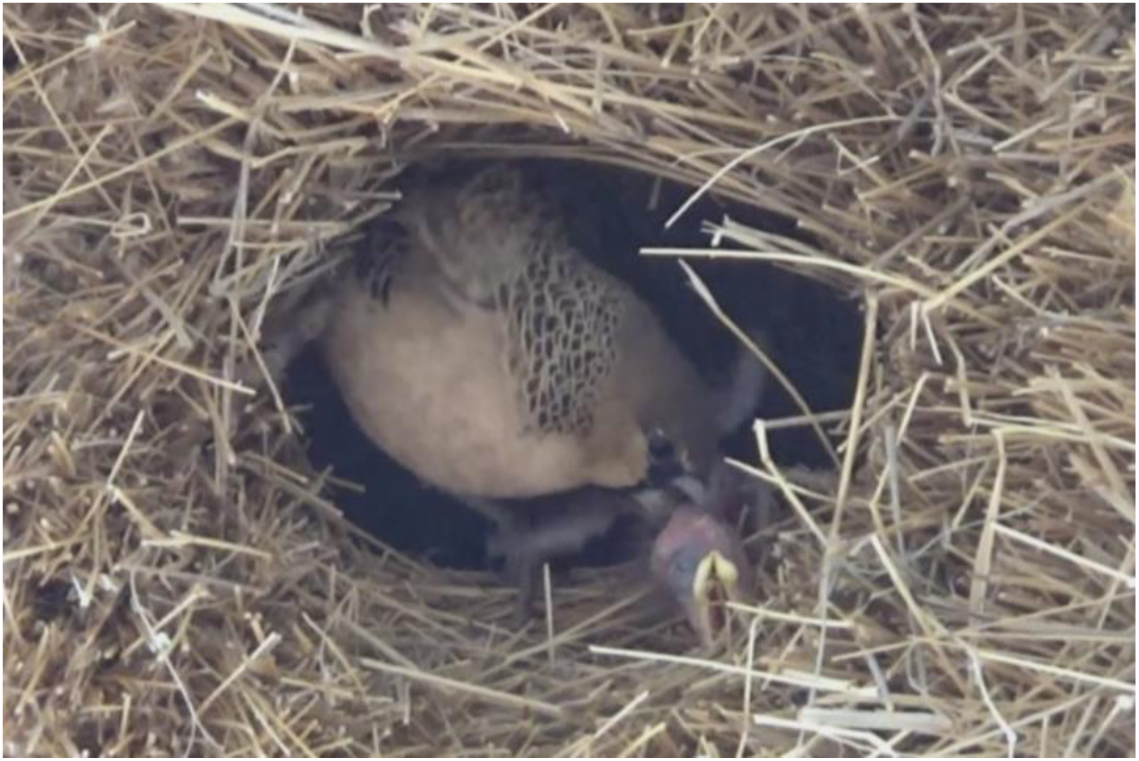
Infanticide event in a sociable weaver nest chamber. An aggressor attacks a nestling and forcefully ejects it from the nest.

If infanticide is motivated by reproductive benefits, it should be associated with increased access to reproduction, *i.e.* it should lead to mating with one of the parents of the young killed to access to a breeding chamber, or both. If the reproductive benefits are associated with competition for food for the developing offspring, we expect that the aggressors will be reproducing at the time of committing infanticide or soon after. Alternatively, if infanticide in sociable weavers is associated with a life-history trade-off, we predict that it should be committed by the parents after suffering partial predation, or when conditions for raising nestlings are poor. Partial nest predation is relatively rare in sociable weavers (the main nest predators are snakes that typically take entire clutches; Covas, Du Plessis, et al., 2008; D’Amelio et al., 2022). However, climatic conditions are extremely variable and associated with offspring mortality (D’Amelio *et al*., 2022), which could lead to situations where raising the brood is no longer viable for parents.

## METHODS

The sociable weaver is a small (ca. 27.5g) sexually monomorphic passerine endemic to the semi-arid Kalahari savannahs of southern Africa (Maclean, 1973a). Both sexes display a black plumage patch that is typically fully developed between 4-5 months of age (Maclean, 1973b). They are obligate colonial, living in colony structures that vary in size from less than 10 to several hundred individuals (Maclean, 1973a). Sociable weavers are facultative cooperative breeders and within each colony there are social clusters comprising breeding pairs and their helpers. The number of pairs with helpers varies between breeding seasons, ranging 30-80% (Covas et al., 2006). Helpers are usually the offspring from previous breeding attempts (from one or both breeders, and can be male and female), but up to 14% of helpers are more distant relatives (e.g., cousins) or unrelated (Covas *et al*., 2006; Ferreira, 2021).

The breeding season usually starts in September or October and continues for as long as the conditions remain suitable, often until April or longer, depending to a large extent on the amount and pattern of rainfall (Mares *et al*., 2017). Females can therefore lay several clutches in the same breeding season (D’Amelio *et al*., 2024). These are hereafter referred to as “breeding attempts”. Clutch size in this species varies between 2-5 eggs (exceptionally 6; Covas et al., 2008; Fortuna et al., 2021; Maclean, 1973b). Incubation lasts 15 days and typically starts on the first or second day of laying, leading to asynchronous hatching. The nestling period usually lasts 21-24 days (Maclean, 1973b).

This study was conducted at Benfontein Nature Reserve, ca. 6 km south-east of Kimberley (Northern Cape Province, South Africa, 28°520S, 24°500E, 1190 m.). Most of the colonies in our study area have been captured and ringed since 1993. Since 2008, all birds in the study colonies have consistently been marked with a unique colour combination during annual captures that usually take place between August-September (before the onset of the breeding season; Covas et al., 2002; Paquet et al., 2015). These captures therefore allow us to identify new (unringed) individuals that have moved into our study population and track movement between colonies. Upon capture, a blood sample is taken from the birds’ brachial vein to genetically determine sex (Griffith *et al*., 1998), parentage and relatedness among individuals (using 16 polymorphic markers; Van Dijk et al., 2015), which allows to differentiate helpers from breeders (see below).

### Breeding monitoring and individual identification

In this study, we use data from the detailed monitoring of breeding activity that was conducted between September 2010 – March 2023 at 13-15 study colonies per year, and videos recorded at these same colonies from 2014-2023. All nest chambers in these colonies are numbered and were inspected every 1-3 days to detect initiation of new clutches and monitor the fate of eggs and nestlings. The day when the first nestling hatches is considered day 1 of the nestling period and these nests are visited again on day 9 (to collect a blood sample weigh and mark each nestling with a metal leg ring) and on day 17 (to weigh, measure and mark nestlings with an individual colour combination). As part of a long-term experiment (Fortuna *et al*., 2021; D’Amelio *et al*., 2022), during this period we have artificially reduced nest predation at 3 to 8 colonies per year by wrapping tree trunks with plastic film. This generally prevented snakes from climbing up trunks and reaching the colonies (see details in D’Amelio et al., 2022). Both protected and natural colonies were included in the present study.

To identify the individuals feeding at each nest, to study nest attendance and determine group size, we collected video-recordings (from 2014 onwards) by placing a camera directly below the nest entrances. Details on the criteria for determining group size and identification of breeders and helpers are given below (see also Fortuna et al., 2021; D’amelio et al., 2024). The video-recordings were conducted in the morning (typically starting 1-3h after sunrise) and recorded both incubation (when mainly the breeding pair visits the nest, as helpers rarely incubate) and the nestling period, when both parents and helpers feed the nestlings. Incubation videos recorded ca. 1.5h (range 1-15 incubation days, 1.94 ± 0.74, mean ± SD hours) and we analysed 1332 videos for 1208 breeding attempts. Provisioning videos were recorded during the nestling period for ca. 4-8h. These videos were predominantly recorded around ages 9 and 17 days, since this follows our long-term protocol (however in this study we used additional recordings available from ages 1-23 days). The number of days recorded per nesting attempt was 2.42 ± 0.87, mean ± SD) and we analysed 5423 videos for 2109 breeding attempts.

### Infanticide recordings

The videos obtained during this period allowed us to record several infanticide attempts and, in some cases, to identify the individuals performing them – hereafter called ‘aggressors’ (see https://doi.org/10.5281/zenodo.18385267 Covas et al., 2026). We considered that there was an infanticide attempt whenever we saw clear aggression of nestlings that were alive (e.g. repeated pecking or unambiguous bloody pecking marks) and/or we saw an attempt to expel live nestlings from inside the nest. If the nestling was not seen moving at any point during the ejection event, we considered that it had died for other reasons and was being removed from the nest for sanitation. Removal of chicks that were already dead inside the nest occurred in 15 cases, which were excluded from our analyses (Fig. ESM1). We also excluded seven cases where nestlings fell off the nest when no adult was present (Fig. ESM1). This could result from pushing by other nestlings or from an accidental fall. Additionally, we recorded ejections of 12 eggs (from seven broods; Fig. ESM1) but these were not considered in this analysis given that it was impossible to know whether the eggs were fertile, and hence we could not apply the same criteria used for the nestling ejections. In addition, there is greater uncertainty about group membership (i.e. future helpers) at the egg stage (but see Fig. ESM1 suggesting that both parents and intruders can remove eggs).

For each infanticide attempt observed, we recorded whether it was successful (i.e. the chick was seen being pushed out of the nest chamber) or non-successful (i.e. the aggressor was interrupted by other birds or the nestling was able to physically resist the ejection).

### Age, sex, and relatedness to the nestlings

Age corresponds to the exact age for all individuals born in the study colonies (where all nestlings have been ringed since 2010). Immigrants were attributed an age of 690 days for immigrant males and 727 days for immigrant females on the date of first capture, which is based on the average known age of first dispersal from our long-term captures data (n= 4441 birds of known age; 9.84 % of these were immigrants that moved within the study area; unpublished data). Sociable weavers present a fully developed adult plumage after ca. 6 months old and from this time their age can no longer be determined by morphological features (R. Covas pers. obs.).

Sex was determined by genetic analyses and the genetic relatedness to nestlings was obtained from a pedigree dataset that resulted from a combination of genetic, breeding history and video analyses (for more details see Fortuna et al., 2021; Silva et al., 2018). These data allowed us to differentiate three categories of individuals: (1) breeders, (2) helpers or (3) intruders. Breeders were individuals that were seen feeding the nestlings and that were assigned as parents through the genetic analyses (or combination of genetic analyses and incubation videos in case of ambiguity). Helpers were individuals that were seen feeding the nestlings at least three times, but were not assigned as parents through the criteria described above. All remaining individuals seen entering the nest are defined as ‘intruders’. In addition, we use the term ‘aggressors’ to refer to any birds that conducted an attack, regardless of whether they were members of the breeding group or intruders. Two individuals (n=4 nestling ejections) were inconclusive in their relationship to the brood (i.e. genetic data was missing and life-history and video data were insufficient to conclude on relatedness to the nestlings) and thus were not considered in these relatedness calculations (but were included in the sex-ratio and age calculations, when these data were available).

### Movement and reproductive history of the aggressors

Colony of origin and residency of the aggressors were the criteria used to determine resident/immigrant status. We considered as resident the birds born at the colony where they committed infanticide.

We used the aggressors’ reproductive history to assign them to one of three breeding categories. 1) Currently breeding, if the individual was known to have a nest with eggs or nestlings at the time of the infanticide attempt; 2) previous breeder, if the bird was known to have been previously reproducing at that same colony and 3) non-breeder, if the bird was never recorded breeding until the date when the infanticide attempt took place.

### Future reproduction and nest occupancy

To understand whether aggressors could increase their access to reproduction because of infanticide, we assessed if they reproduced after the attack, and further scrutinised their future partner acquisition and nest occupancy. Only birds that were ringed with full colour combinations were used for this screening. For partner acquisition, we determined if aggressors subsequently reproduced over the next 365 days, and if they did, whether they paired with one of the breeders of the attacked nest. For nest occupancy, we determined if aggressors subsequently reproduced in the chamber that was attacked or a nearby chamber (since sociable weavers breeding groups can occupy up to 3 nearby chambers; Covas R., Doutrelant C., Silva L.R., Ferreira A.C. and Fortuna R. unpublished data).

All statistical analyses and calculations were conducted in R and the respective code is available as an electronic appendix and archived on Zenodo (Covas et al., 2026 https://doi.org/10.5281/zenodo.18385267).

## RESULTS

We recorded on video a total of 50 infanticide attempts of nestlings (out of a total of 3095 broods recorded over the course of 9 years). However, we note that these videos were mostly recorded around age 9 and 17 of the nestling period, as per our long-term protocol or breeding monitoring, but infanticide appears to be more common before the bulk of our recordings were conducted (see Fig. ESM2 for the distribution of video effort in relation to nestlings’ age). The infanticide attempts recorded here were conducted at 34 nest-chambers in 11 colonies. One of these attempts was committed on the same nestling simultaneously by two intruders, which means that there were 51 individual attackers recorded for these 50 infanticide attempts. Of the 50 infanticide attempts recorded, 66% (33 nestlings) ended in a successful infanticide (i.e. the aggressors successfully expelled the nestling out of the chamber). In the remainder, the victim’s breeding group members chased the aggressors, or the nestling resisted eviction. The latter was often the case with older nestlings (>9 days old, but our biased sampling in relation to age prevents us from drawing conclusions about any effects of nestling age on the success of the attack; see Fig. ESM2). The average duration of the successful events was 2.91±4.01 minutes (min 0.08, max 17.43, from the moment an individual enters the nest until it concluded the ejection, i.e. one or more nestlings were ejected from the nest; ESM video).

### Sex and age of the aggressors

We were able to identify individually 40 aggressors, but only 27 had known sex. Of these, 59.2% were females (*n*=16) and 40.7% were males (*n*=11), including one instance where two birds jointly killed the chicks of a nest (Fig. 2). However, if we consider the sex-ratio of unringed immigrants that join our population every year (70% of which were females, based on *n*= 416 immigrant individuals captured from 2014-2023 in the same colonies, and which were subsequently sexed), we would estimate that ca. 71% of the aggressors were females.

**Figure 2:**
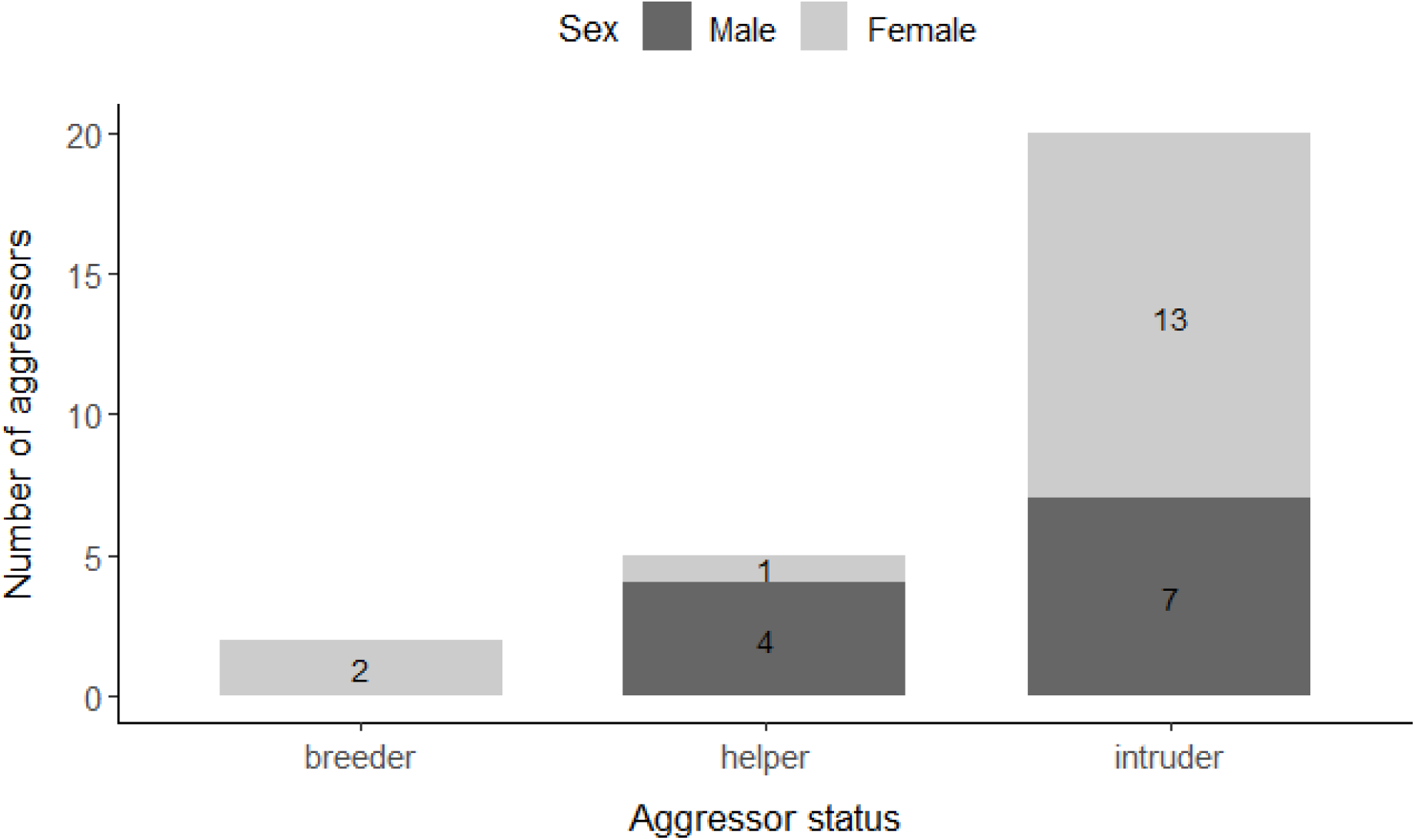
Number of aggressors with confirmed sex information that attempted infanticide (*n*= 27) categorized by sex and status (breeder, helper, intruder). Aggressor sex is indicated by colour: males in dark grey), females in light grey.

All identified aggressors were adult birds. Based on a smaller sample of aggressors and breeders of know age, 79.1% of aggressors were younger than the same sex breeder of the brood they had attacked (*n*= 24 aggressor-breeder dyads; Fig 3).

**Figure 3:**
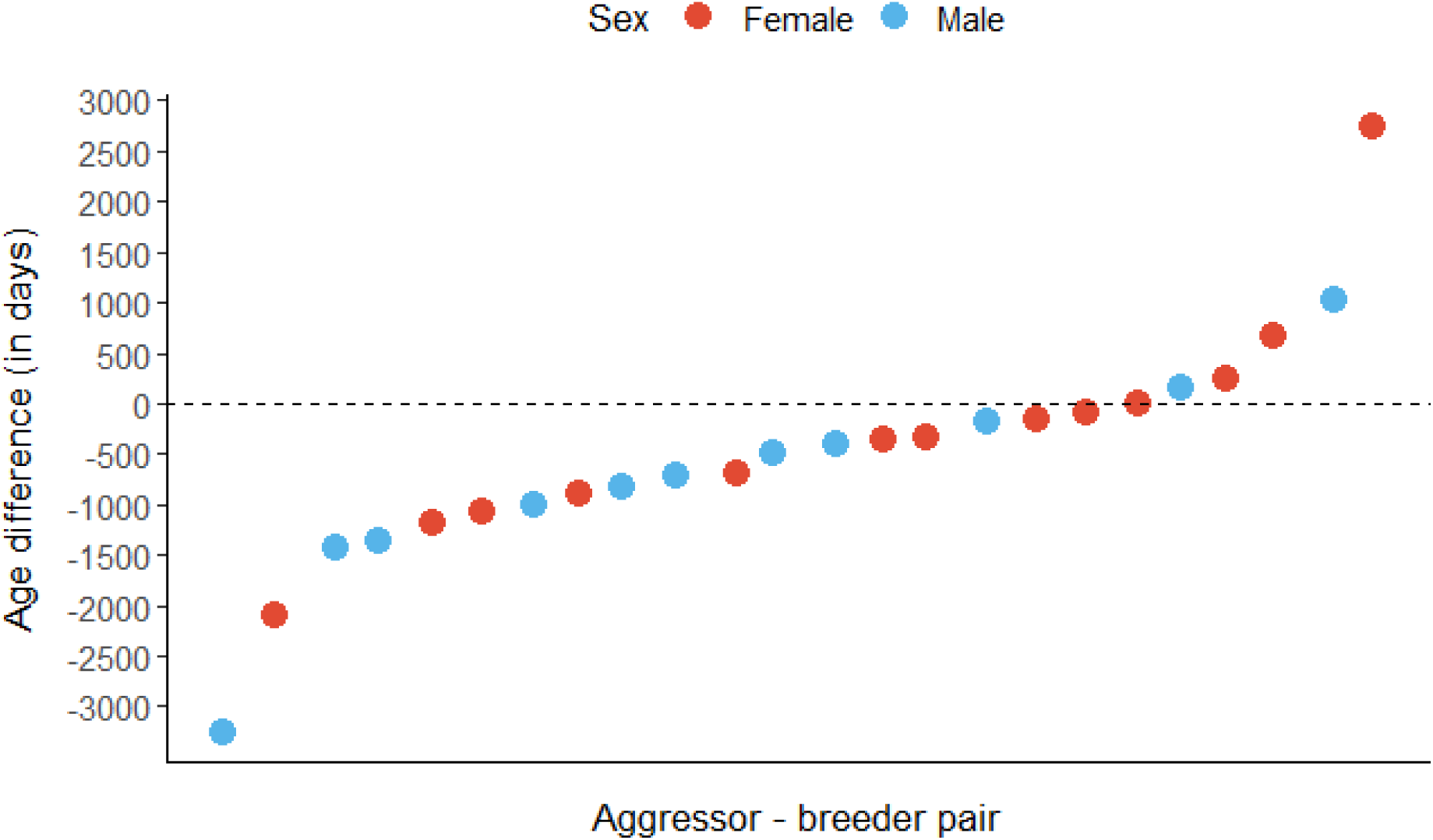
Age difference in days between the aggressors and breeders of known age (24 pairs where 79.1% of aggressors were younger than the same-sex breeder of the brood that they attacked). The age difference was calculated as the age of the aggressor minus the age of the same-sex breeder of the attacked brood. Negative values mean that the aggressor is younger than the same sex-breeder of the focal nest (with differences >365 days in the large majority of cases). The aggressors’ (and correspondent breeder’) sex is represented in red and blue for females and males, respectively.

### Relationship to the nestlings

Aggressors were usually individuals external to the breeding group. We found that 82.5% (of the 40 aggressors identified) were not members of the breeding group (*i.e.* breeders nor helpers). There was, however, evidence of two breeding females ejecting a live nestling from their own nests (5%) and five cases of helpers ejecting nestlings of the group that they were helping (12.5%). Four of these were male helpers, with various degrees of relatedness to the attacked nestling: brother, uncle and great-uncle, and a “social half-brother” that was born as part of a cross-fostering experiment and helped its social parents (*i.e.* it was not genetically related to the nestling ejected). The only female helper that committed infanticide was not related to the attacked nestling but was born in that colony.

### Movement and reproductive history of aggressors

The majority of aggressors were born outside the colony where they committed infanticide (62.5%, *n*= 25 aggressors; Fig. 4). The remainder 37.5% were born at the colony (*n*= 15 aggressors), including the two breeders and the five helpers (Fig. 4). Those that were not part of the breeding group (*i.e.* those considered “intruders”) were mostly born outside of the colony (75.7%, *n*= 25 aggressors). All non-ringed individuals were considered as not born at the colonies of the attacks since all the nestlings born in the studied colonies and all adults captured there were ringed.

**Figure 4:**
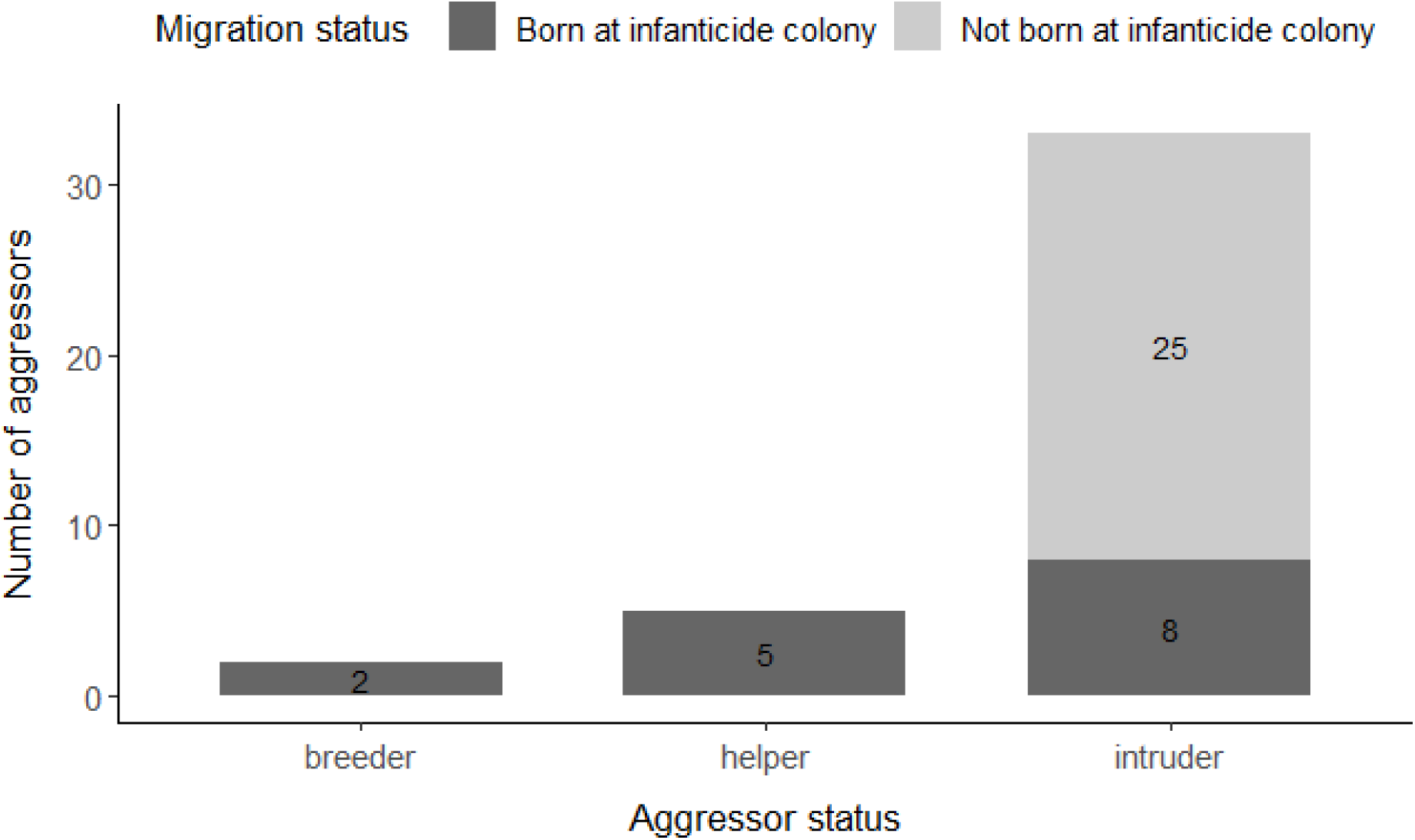
Number of aggressors in relation to their migration status (N=40 aggressors) and their relationship to the nestlings attacked (N= 50 nestlings; aggressors could attack more than one nestling). Aggressor status was defined as breeder, helper and intruder. Migration status was defined as being born (in dark grey) or not (in light grey) at the colony where the attack took place.

The reproductive status of the aggressors at the moment of the attack was unknown in 32.5% of cases, as the birds were not ringed (*n*= 13 assumed different aggressors, which were all intruders). For the birds with known histories, we found that 88.9% (*n*= 24) of the aggressors were not breeding at the time of the attack (Fig. 5), although five were helpers at other nests (or had recent helping activity in the 15 days before or after the attack; one of these birds was a helper at another colony). Among these non-breeding intruders, 54.2% (*n*= 13) were never seen reproducing before committing infanticide, while the rest had previously attempted to breed (45.8%, *n*= 11, all at the same colony of the attack).

**Figure 5:**
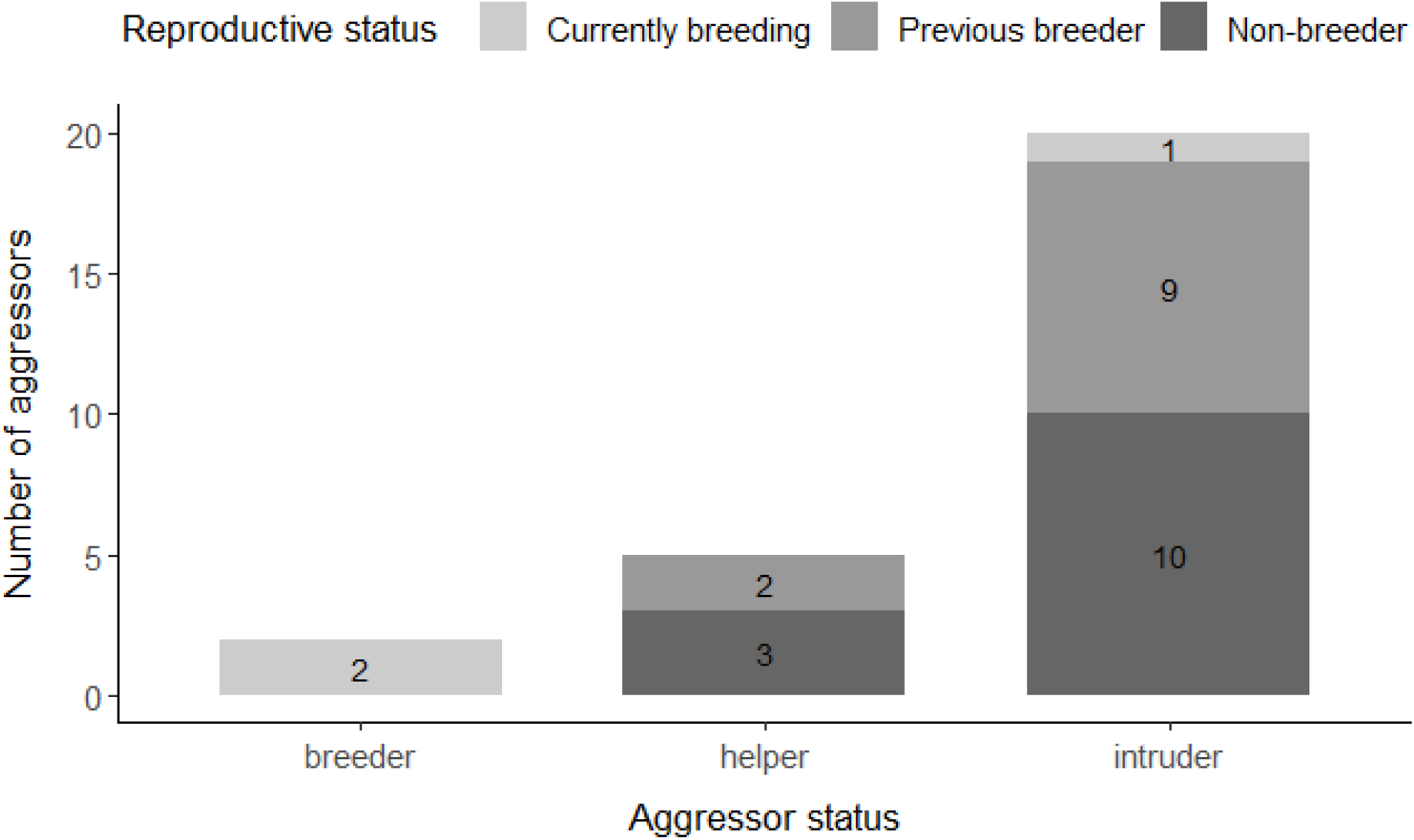
Number of birds that attempted infanticide across different aggressor status categories (breeder, helper and intruder; *n*=27 aggressors). Reproductive status categories are: 1) “currently breeding”, if the individual was known to have a nest with eggs or nestlings at the time of the infanticide attempt (light grey); 2) “previous breeder”, if the bird was known to have been previously reproducing at the same colony (intermediate grey) and 3) “non-breeder”, if the bird was never recorded breeding until the date when the infanticide attempt took place (dark grey).

### Future reproductive activity

With the exception of the two breeders that committed infanticide, we could only identify the subsequent activity of 57.9% (*n*= 22) of aggressors (i.e. birds of known identity and known to be present in monitored colonies after the infanticide). Of these, only 59.1% (*n*= 13; 8 females and 5 males) were observed reproducing 365 days after the infanticide (this corresponds to 2 birds that were helpers and 11 that were intruders when they committed infanticide). For these birds, those reproductive attempts corresponded to their first in 38.5% of cases (n=5). Finally, the majority of these birds reproduced in the same colony where they conducted the attack (84.6%; *n*= 11).

None of these aggressors was recorded reproducing with one of the breeders or helpers (female or male) from the victims’ nest up to 365 days after the attack (i.e. encompassing a full breeding season the following year). Similarly, none of the 22 known aggressors was found to gain access to the chamber attacked or to a nearby (adjacent) chamber over the following breeding season.

## DISCUSSION

Infanticide is challenging to study as it is unpredictable in occurrence and can be very brief (average time was less than 3 min in this study). Our long-term study has enabled us to gather multiple observations across the years, providing unambiguous evidence that infanticide behaviour occurs in sociable weavers at the nestling stage (and also at the egg stage; Fig. ESM1). Over 80% of the infanticide attempts considered here, were conducted by birds that were not part of the breeding group, and over 60% of these birds were born in a different colony. In addition, aggressors were usually younger than the parents of the victims. Aggressors were usually not reproducing when they committed the attacks. No evidence was found that the aggressors subsequently paired to one of the breeders or helpers from the nest attacked, or that they occupied the chamber from where the chicks were evicted or a nearby chamber. Of the identified aggressors, just over half were found as breeders 365 days after the attack, most of these reproduced at the same colony, some of them for the first time.

These findings allow us to exclude several mechanisms as possible benefits of infanticide behaviour in this species. First, given that the large majority of birds committing infanticide were not the parents or other members of the breeding group, there were few cases where infanticide of nestlings in sociable weavers could be motivated by a life-history trade-off. Convincing data of life-history trade-offs motivating infanticide in birds was found in *Eclectus* parrots, where sons in poor-quality nest hollows were killed (Heinsohn *et al*., 2011), and in grey-capped social weavers where infanticide was associated with partial nest predation, making it more advantageous to eject the whole clutch and start again, than to raise a reduced clutch (Cheng *et al*., 2019). In our study, we had no information about the condition of the chicks that were ejected by their own mother, and they could have been sick or in poor condition. In one of the cases, the other nestlings from the brood were all ejected; in the other case, the three other siblings in the nest fledged. Hence, it is possible that a life-history trade-off explains some cases of infanticide in sociable weavers, but it is not possible to draw any conclusions based on these two cases, and this mechanism cannot account for the large majority of cases of infanticide behaviour in this species.

Second, our results do not provide any evidence of increased access to a mate or nesting chamber from the group which suffered infanticide. Here, none of the birds committing infanticide was observed to later reproduce with a parent or helper of one of the victims over the following year. We also found no evidence that the aggressors gained access to a nesting chamber, which is an important asset in this species since it is used for reproduction, roosting and shelter throughout the year (Maclean, 1973a). Access to a chamber could therefore have improved the probability of breeding for these aggressors. Finally, with the exception of the two breeding females, only one aggressor (a male) was reproducing when committing infanticide, suggesting that direct competition for feeding offspring was not a main factor motivating the majority of the attacks. Hence, our results contrast with studies conducted in mammals, which suggest that infanticide can lead to increased chances of reproduction or to a larger share of resources for the offspring of individuals that are breeding simultaneously (Lukas and Huchard, 2014, 2019).

Of the identified aggressors that could be tracked after the attacks, the majority were born in another colony, and hence had no relatives in the colony where they committed the attacks, but became breeders there at a later stage. It might be possible that reducing the reproductive success of other colony members could contribute to reduce competition for resources for future offspring, and directly for the aggressors, favouring the aggressor’s genetic lineage over others in the colony. Although we are not aware of empirical studies demonstrating such a mechanism (but see Hrdy, 1979), we speculate that this could take place in two ways. First, given that sociable weavers are cooperative breeders and the helpers are most commonly the offspring of one or both breeders (Covas et al., 2006; Ferreira, 2021), current reproductive success will influence the future probability of having helpers at the nest. A large group with more helpers is likely to be a stronger competitor within a colony than a single pair or a small group. Helpers contribute to groups tasks such as nest maintenance and defence (Ferreira, 2015; Silva, Covas and Doutrelant pers obs) and larger groups occupy more chambers within the colony (Silva, Covas and Doutrelant pers obs) and have higher reproductive success (D’Amelio *et al*., 2022b). Hence, infanticide could contribute indirectly to increase the aggressors’ future reproductive output or to benefit close relatives through improved access to resources and a more favourable social environment for their future offspring, i.e. it could be a form of spite (West and Gardner, 2010). Studies in some group-living species showing that groups can kidnap the offspring of others (Ridley *et al*., 2022) suggest that competition for larger group sizes can be important in animal societies (Heinsohn, 1991; Ridley *et al*., 2022).

We also found a tendency for the attackers to be younger than the breeders, which is compatible with the hypothesis of competition among family groups within the colonies, as newcomers could attempt to increase their status or (future) share of resources in relation to other groups. This, would not be the case of the five helpers that committed infanticide at the nest that they were helping, as four of these were males that were related to the nestlings attacked (the kinship ranged from brother to great uncle). However, two reproduced at the colony after the attacks. Increased infanticide by younger group members was also found in free-ranging house mice *Mus musculus*, where virgin females were more likely to commit infanticide than pre-pubertal ones (Mccarthy and Saal, 1985). However, in the house mouse, it is not known whether this age-effect on infanticide behaviour is adaptive. Future work will need to examine the trajectories of the pairs and families that have committed and suffered infanticide in sociable weavers to better evaluate whether competition among groups or clans is a possible explanation for infanticide in this species. Specifically, it is possible to investigate whether the subsequent reproductive success or group size of aggressors differs from that of immigrants and other individuals that have not committed infanticide (although this requires an important field effort to ascertain who has not committed infanticide) or is higher than the victims’.

Another non-exclusive explanation is that infanticide could arise from direct competition for food around the colony, as a way of reducing the numbers of competitors (see also Hrdy, 1979). A nest predator exclusion experiment, ongoing at some of our colonies, will allow us to address this question in the future, as reproductive success is higher at these colonies, leading to higher colony density (D’Amelio *et al*., 2022a), which should increase competition for resources. In addition, we have found previous evidence that in sociable weavers, the number of nest ejections increases under hot and dry conditions (D’Amelio *et al*., 2022a), when food resources are presumably scarce. In that study, it was not possible to determine which proportion of these ejections were victims of infanticide or were already dead in the nest. An increase in infanticide under hot and dry conditions could also be a direct consequence of parents and helpers spending more time away from the nests in order to find food, thereby leaving the nestlings alone for longer. Examining the link between hot and dry conditions and infanticide is particularly relevant, in this and other species, given that the frequency of hot and dry periods is increasing as a result of global climate change.

Finally, our data suggest that infanticide behaviour is also likely to be associated with costs. We found that successfully committing infanticide takes effort, with an average of nearly three minutes per successful infanticide attempt, and our videos show that the aggressors physically wrest with the nestlings in order to evict them (ESM video). In addition, both the breeding pair and their helpers fight intruders that attempt to enter the chamber with physical attacks and displacements. Hence, committing infanticide appears to be physically demanding, aggressors undertake the risk of being pursued, and even possibly injured. Other potential costs are likely to involve social retaliation, or a decrease in colony size (since sociable weavers’ survival tends to be higher in larger colonies; (Brown *et al*., 2003). A better understanding of the costs of infanticide is important in order to establish whether infanticide is an adaptive behaviour, since costly behaviours are only expected to evolve if the benefits outweigh the costs.

In conclusion, this study provided clear evidence of infanticide in a highly social and cooperative bird, and provided results that are not compatible with prevalent hypotheses suggesting that it provides reproductive benefits, such as increased access to breeding partners, nest-chambers or a greater share of resources for offspring. We found that there could be different, non-exclusive explanations for infanticide. Life-history trade-offs might motivate infanticide in this species, but on rare occasions. Competition among breeding groups or clans and indirect reproductive benefits are potential mechanisms that are compatible with the majority of the cases presented here. However, future work following individuals and groups over extended periods and examining possible interaction with climatic conditions are required to establish the benefits – and costs – of infanticide in this species, and whether it is promoted by specific environmental conditions.

## Acknowledgements

We are very grateful to the field managers, assistants and student involved in the long-term data collection, especially for help with the yearly captures and breeding monitoring. We thank Franck Theron for his work in the field, training and coordination of the field team and management of the sociable weaver long-term database. De Beers Mining Corporation provided access to Benfontein Nature Reserve and logistical assistance. The sexing and genotyping analyses were conducted at the CTM lab, CIBIO, and we thank Susana Lopes and all the technical staff that conduct the work.

## Data, scripts, code, and supplementary information availability

Supplementary figures and texts are available on BiorXiv (Covas et al. 2025, https://doi.org/10.1101/2025.05.27.656097). The datasets, code and auxiliary materials are archived on Zenodo (DOI: https://doi.org/10.5281/zenodo.18385267).

## Conflict of interest disclosure

The authors declare that they comply with the PCI rule of having no financial conflicts of interest in relation to the content of the article. M. Paquet is a recommender and data editor at PCI.

## Funding

The study was funded by FCT (Portugal, grants IF/01411/2014/CP1256/CT0007 and PTDC/BIA-EVF/5249/2014), ERC (EU, Consolidator grant 866489) and the DST-NRF Centre of Excellence at the Fitzpatrick Institute of African Ornithology University of Cape Town (South Africa) to R.C and additional funding from ANR (France, grants 15-CE32-0012-02 and 19-CE02-0014-01) to C.D.. A.C.F was funded by University of Zurich Forschungskredit postdoc grant (K-74312-01-01 University of Zurich), Swiss Federal Commission for Scholarships and by ERC grants 866489 to R.C and 850859 awarded to Damien Farine, R.C. was funded by FCT (IF/01411/2014/CP1256/CT0007 and CEECIND/03451/2018). PBDA was supported by a South African Claude Leon Postdoctoral Fellowship 2019-20, a Marie Curie Individual Fellowship “SOCIAL MATCH” N. 896475 and by ANR 19CE02-0014-01. R.F. was funded by FCT (SFRH/BD/130134/2017 and COVID/BD/151899/2022 https://doi.org/10.54499/COVID/BD/151899/2022). The Sociable Weaver Project is supported by the French OSU-OREME, and the CNRS programme on Long-Term Studies in Ecology and Evolution (SEE-Life).

## Notes

### Competing Interest Statement

The authors have declared no competing interest.

### Summary of Updates

This version has minor edits and modifications for increased clarity following comments by Data Editors and a Recommender from PCI. This is the final version as recommended by PCI

